# Axis formation in annual killifish: Nodal coordinates morphogenesis in absence of Huluwa prepatterning

**DOI:** 10.1101/2021.04.16.440199

**Authors:** Philip B. Abitua, Deniz C. Aksel, Alexander F. Schier

## Abstract

Axis formation in fish and amphibians is initiated by a prepattern of maternal gene products in the blastula. The embryogenesis of annual killifish challenges prepatterning models because blastomeres disperse and then re-aggregate to form the germ layers and body axes. This dispersion-aggregation process prompts the question how axis determinants such as Huluwa and germ layer inducers such as Nodal function in annual killifish. Here we show in *Nothobranchius furzeri* that *huluwa*, the factor thought to break symmetry by stabilizing β-catenin, is a non-functional pseudogene. Nuclear β-catenin is not selectively stabilized on one side of the blastula but accumulates in cells forming the incipient aggregate. Inhibition of Nodal signaling blocks aggregation and disrupts coordinated cell migration, establishing a novel role for this signaling pathway. These results reveal a surprising departure from classic mechanisms of axis formation: canonical Huluwa-mediated prepatterning is dispensable and Nodal coordinates morphogenesis.

**One Sentence Summary:** Axis formation in annual killifish relies on Nodal to coordinate cell migration and is independent of Huluwa-mediated prepatterning.

## Main Text

Spemann and Mangold discovered that cells on the dorsal side of gastrulating newt embryos were capable of inducing a secondary axis when transplanted into the ventral side of a host embryo (*1*). They named this group of cells the ‘organizer’ for its ability to organize surrounding tissue into a separate body axis. It is believed that anamniotes (i.e. fishes and amphibians) generate the organizer by breaking radial symmetry via a prepattern of maternally provided mRNAs. In particular, the maternal determinant *huluwa* is partitioned to the future dorsal side, culminating in the stabilization and nuclear accumulation of β-catenin, which acts as a transcriptional activator of organizer-specific genes (*2*). After dorsal-ventral axis determination, the TGF-β morphogen Nodal induces the phosphorylation of the downstream effector Smad2 in target cells, resulting in mesoderm and endoderm specification (*3*).

Among teleosts, annual killifish have an atypical development that challenges current models of axis formation (*4*). While the cells of the extra-embryonic enveloping layer (EVL) expand over the surface of the yolk as in a conventional epiboly, the deep blastomeres completely disperse and only re-aggregate later to form the embryo proper (Fig. 1A). At the onset of epiboly these blastomeres de-adhere from one another and spread across the expanding EVL through ‘contact inhibition of locomotion’, a phenomenon where cells change their direction upon collision with one another (*5, 6*). At the end of epiboly, the deep cells of annual killifish are completely dispersed as single cells under the EVL (*4*). In contrast, blastomeres in other teleosts, such as zebrafish and medaka, stay adherent and intercalate (*3, 7*). The dispersion and randomization of deep blastomeres raises the question of whether prepatterning is involved in axis specification and how the inducers Huluwa and Nodal act in annual killifish. To address these questions, we chose the annual killifish *Nothobranchius furzeri*, owing to its genetic tractability and availability of a reference genome (*8, 9*).

**Figure 1.**
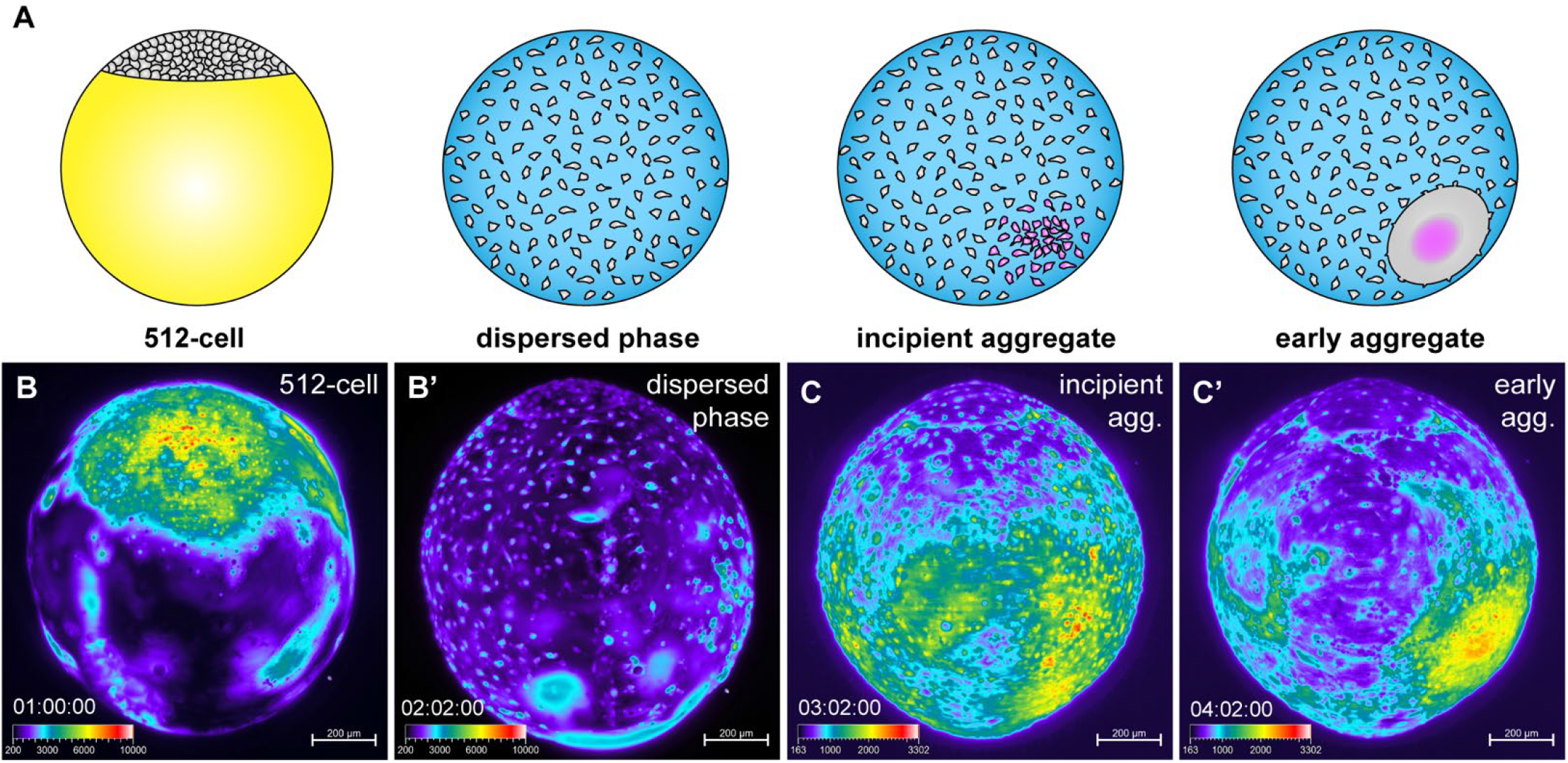
Nuclear β-catenin dynamics during early *N. furzeri* development. (**A**) Schematic of *N. furzeri* stages. Yellow represents the yolk not yet enveloped by the EVL and magenta represents the mesoderm. (**B**) Snapshots from a time-lapse of transgenic BC1:egfp embryo starting at approximately the 512-cell stage during the onset of dispersion. (**B’**) View at the end of epiboly. (**C**) Snapshots from a time-lapse of transgenic BC1:egfp embryo starting at the incipient aggregate stage. (**C’**) View during early aggregate stage, 24 hours later. All images are lateral views. Timestamp indicates days: hours: minutes starting from fertilization. The signal intensity is color coded (see intensity scale).

To understand how the embryonic axes form in *N. furzeri*, we analyzed the expression of marker genes and studied the activity of the Huluwa/β-catenin and Nodal pathways. We first sought to visualize when and where β-catenin was stabilized during *N. furzeri* development. Using a β-catenin antibody against zebrafish Ctnnb1, we observed uniform β-catenin nuclear localization prior to dispersion (256-cell stage) (fig. S1A), in contrast to the asymmetric localization observed in other anamniotes (*10, 11*). During the dispersed phase, β-catenin staining was markedly reduced (fig. S1B). Nuclear β-catenin accumulated again in a cluster of cells that formed the incipient aggregate (fig. S1C). As re-aggregation continued, nuclear β-catenin was largely restricted to peripheral cells that were in the process of joining the aggregate (fig. S1D, above the dotted line). To directly monitor β-catenin dynamics in live embryos, we generated a transgenic line that expressed a fluorescently tagged nanobody that binds to β-catenin (*12*). Live imaging showed a decrease in β-catenin levels as cells dispersed, consistent with the observations made with the β-catenin antibody (Fig. 1B, fig. S1B, and movie S1). During incipient re-aggregation, β-catenin was seen accumulating in the nuclei of a cluster of individual cells that subsequently formed the center of the future aggregate (Fig. 1C, fig. S1C, and movie S2). Remarkably, the accumulation of nuclear β-catenin was dynamic, as cells near the future aggregate only transiently accumulated nuclear signal (fig. S2, and movie S2). Eventually, the nuclear β-catenin signal stabilized as re-aggregation continued, and cells adhered to each other (Fig. 1C’, and fig. S2). The absence of a stable pattern before the formation of the incipient aggregate suggests that β-catenin does not act as a maternal dorsal determinant in the early blastula of *N. furzeri*. Instead, its later accumulation marks the first site of aggregation.

We next asked whether Nodal signaling also differs between *N. furzeri* and other anamniotes. In zebrafish, Nodal gene expression and pathway activity are found at the blastula margin and then become restricted to the axial mesoderm (*13*). By contrast, the *N. furzeri* Nodal gene *ndr2* showed broad and weak early expression in deep cells during the dispersed phase (Fig. 2A). At the incipient aggregate stage, *ndr2* expression was enriched in a cluster of cells where cell-cell adhesion had initiated (Fig. 2B). The timing and location of *ndr2* expression correlated with the emergence of nuclear β-catenin in the incipient aggregate (Fig. 1C). As cells continued to join the early aggregate, *ndr2* expression and nuclear Smad2 staining also was strongest in the aggregate center (Fig. 2, C and D). *Lefty1*, a Nodal feedback inhibitor (*3*) was also expressed in a cluster of deep cells at the site of the incipient aggregate (Fig. 2E). Thus, in contrast to zebrafish, *N. furzeri* Nodal signaling is initially broadly active and then becomes restricted to the center of the aggregate.

**Figure 2.**
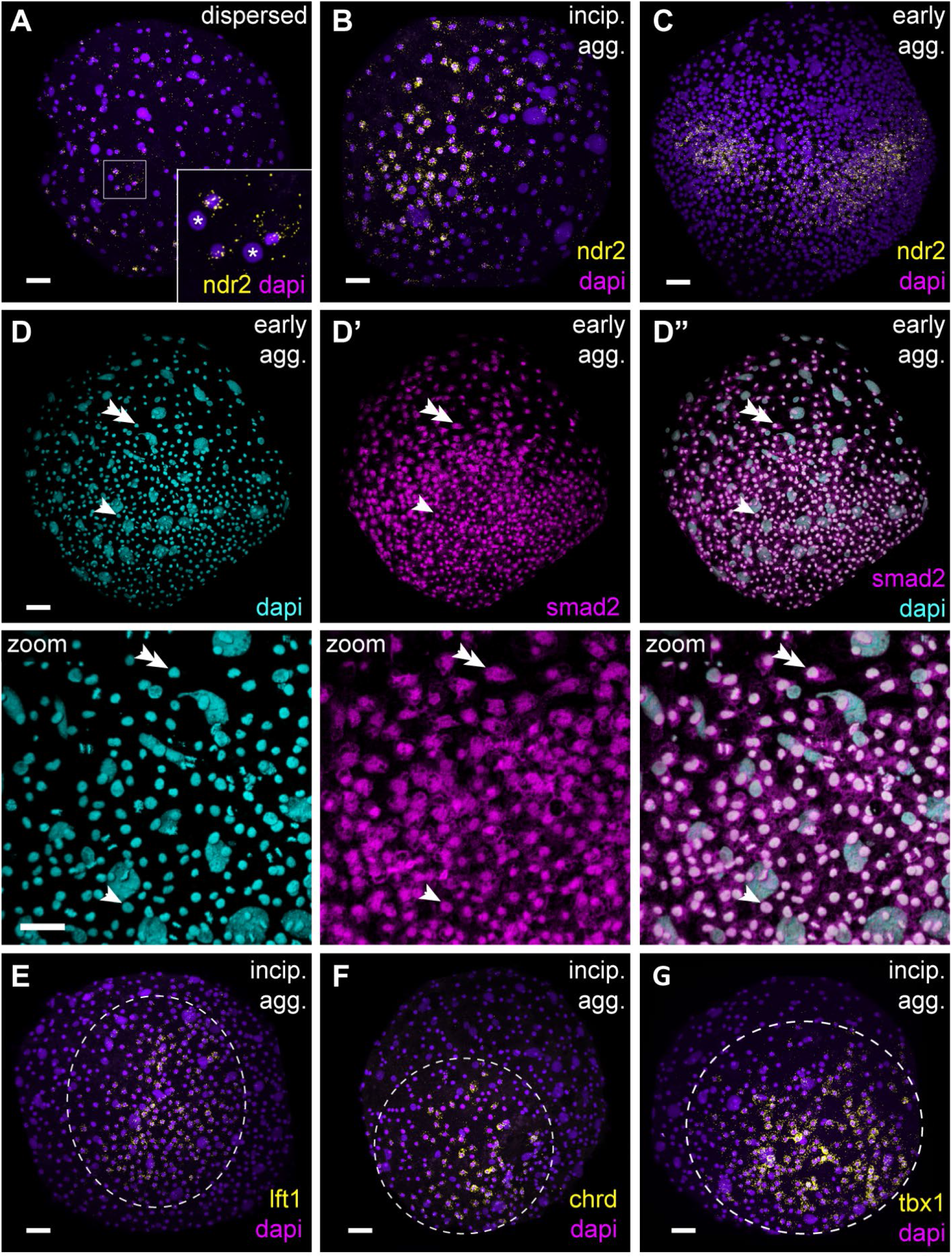
Pattern formation during *N. furzeri* development. (**A**) *Ndr2* mRNA expression during the dispersed phase. Inset shows zoom-in of boxed area. Asterisks indicate EVL cells that lack *ndr2* mRNA. (**B**) *Ndr2* mRNA expression during the incipient aggregate stage. (**C**) *Ndr2* mRNA expression during the early aggregate stage. (**D**) Smad2 shows nuclear localization closer to the center of re-aggregated deep cells (compare arrowhead to double arrowhead). Zoom-in view shows the periphery of the aggregate, which is positioned at the bottom. (**E**) *Lefty1* mRNA expression, (**F**) *Chordin* mRNA expression, and (**G**) *Tbx1* mRNA expression during the incipient aggregate stage. Dotted line demarcates the area of expression in (E) to (G). (A) to (C) and (E) to (G) aggregate views; (B) lateral aggregate view. Scale bars, 50 μm.

The emergence of pattern during the incipient aggregate stage is also reflected in the expression of other conserved germ layer marker genes. We observed the expression of *chordin*, a BMP antagonist secreted from the dorsal organizer of vertebrates, in the incipient aggregate (Fig. 2F). This site of the future aggregate also expressed the mesodermal marker gene *tbx1* (Fig. 2G). Together, these results indicate that the incipient aggregate of *N. furzeri* corresponds to the dorsal organizer and presumptive mesoderm and is the site of high β-catenin and Nodal signaling activity.

As development proceeds, germ layer marker gene expression becomes more similar to other teleosts. Future ectodermal cells joined the aggregate and surrounded the presumptive mesoderm, which then internalized to define the body axes (fig. S3). At the bud stage, *lefty1* was expressed throughout the entire axial mesoderm (fig. S3A). *Chordin* was also expressed in the axial mesoderm but was downregulated in the posterior most region where *tbx1* was expressed (fig. S3, B and C). The ectodermal marker *sox2* was expressed in the surface ectoderm and was absent from the axial mesoderm (fig. S3D). The midbrain-hindbrain boundary was demarcated by the expression of *otx2* and *gbx1* (fig. S3, E and F). Despite the unusual early morphogenesis where deep cells dispersed, migrated and then re-aggregated, by the bud stage, *N. furzeri* expression patterns were typical for teleosts, including for a related annual killifish of the genus *Austrolebias* (*6, 13–15*).

The absence of a clear pattern prior to re-aggregation raised the question what role *huluwa* might have in *N. furzeri* axis formation. We identified the *N. furzeri huluwa* orthologue on chromosome Sgr02. Strikingly, aligning the genomic sequence of *N. furzeri huluwa* with other teleosts, including the closely related non-annual mangrove killifish, *Kryptolebias marmoratus*, revealed a single base pair insertion that results in a predicted frameshift mutation (Fig. 3A). Alignment of *N. furzeri huluwa* mRNA with other *huluwa* orthologues revealed that the predicted *N. furzeri* sequence was truncated, lacking the conserved and essential C-terminal intracellular domain (Fig. 3, B and C) (*2*). These findings indicate that *N. furzeri huluwa* degenerated into a pseudogene.

**Figure 3.**
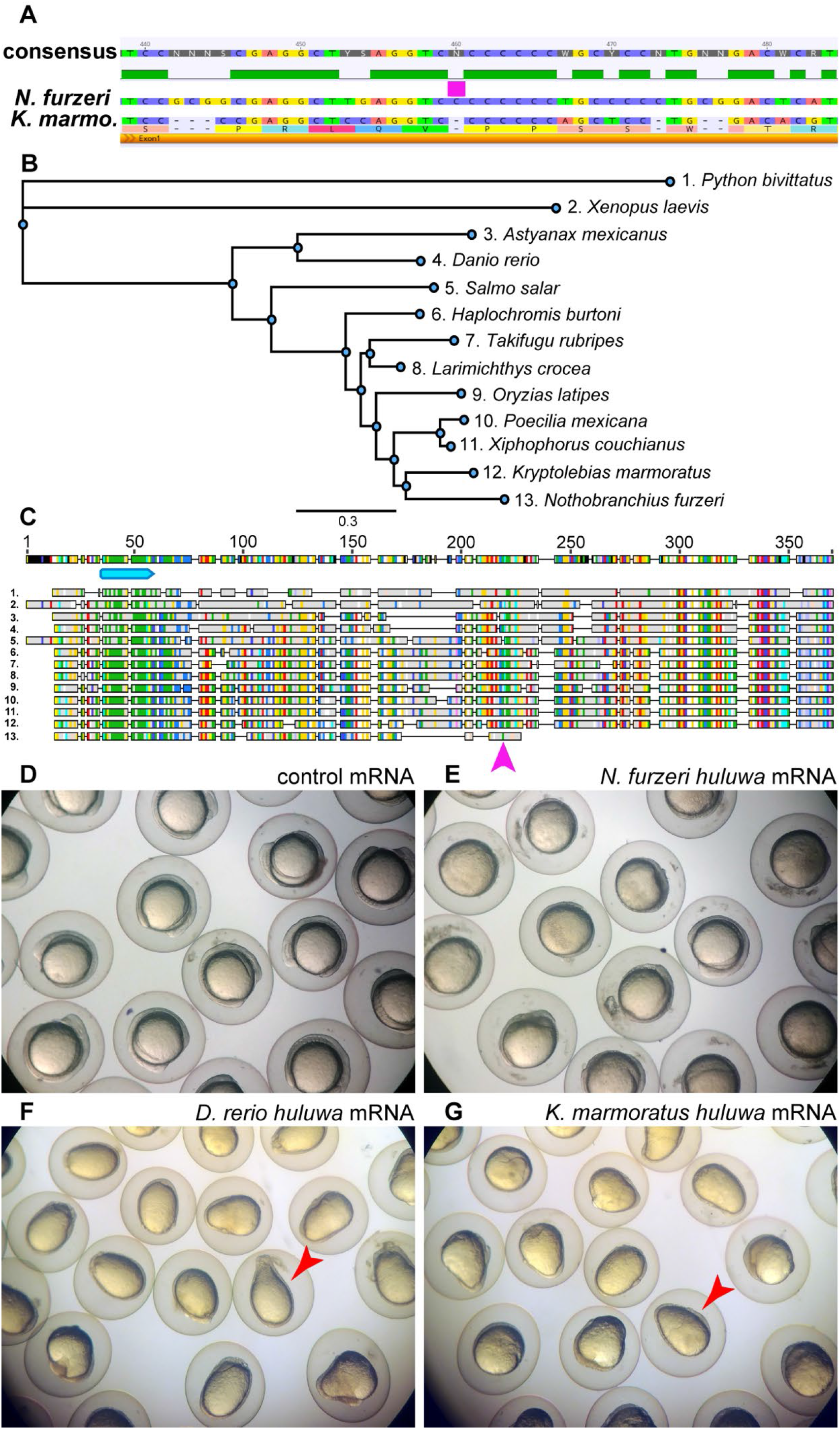
*N. furzeri huluwa* pseudogene lacks dorsalizing activity. (**A**) Alignment of genomic sequences for *N. furzeri* and *K. marmoratus huluwa*. Magenta bar indicates the single base pair insertion in the *N. furzeri* sequence. (**B**) Phylogenetic tree of Huluwa orthologous proteins in 13 anamniote species. (**C**) Alignment of Huluwa orthologous proteins. Numbers correspond to the species indicated in (B). Amino acids are colored using the RasMol scheme. Cyan bar indicates the conserved transmembrane domain. Magenta arrow indicates the site of the frameshift mutation in the *N. furzeri huluwa* sequence (#13). (**D**) Embryos injected with 50 pg of control mRNA at the 1-cell stage (0% severely dorsalized, n = 91). (**E**) Embryos injected with 50 pg of *N. furzeri huluwa* mRNA at the 1-cell stage (0% severely dorsalized, n = 84). (**F**) Embryos injected with 50 pg of D. rerio *huluwa* mRNA at the 1-cell stage (94.6% severely dorsalized, n = 92). (**G**) Embryos injected with 50 pg of *K. marmoratus huluwa* mRNA at the 1-cell stage (95.4% severely dorsalized, n = 107). Red arrowheads in (F) and (G) indicate embryos with an ovoidal morphology, indicative of severe dorsalization. Images of embryos in (D) to (G) were acquired at the 6-somite stage.

To directly compare the activities of *N. furzeri huluwa* to its orthologues, we misexpressed *huluwa* from *N. furzeri*, zebrafish and *K. marmoratus* by mRNA injection into 1-cell zebrafish zygotes. Embryos misexpressing zebrafish and *K. marmoratus huluwa* had the ovoidal morphology of dorsalized embryos (*16*) and died during segmentation (Fig. 3, D to G). In striking contrast, *N. furzeri huluwa* induced cell division defects but had no dorsalizing activity (Fig. 3E). These results reveal that *N. furzeri huluwa* is non-functional as a dorsalizing determinant.

The surprising observation that Huluwa is not conserved as an early dorsal determinant in *N. furzeri* prompted us to test the conserved roles of Nodal signaling in mesendoderm induction. We incubated dispersion phase embryos in A-83-01, a selective inhibitor of Activin/Nodal type 1 receptors (*17*), or injected mRNA for the Nodal inhibitor *lefty1* at the 1-cell stage. Nodal-inhibited embryos failed to express *ndr2, lefty1, chordin*, and *tbx1* (Fig. S4), consistent with the role of Nodal signaling in mesoderm induction. Initial cleavage divisions and epiboly were not affected (Fig. 4, A to C), but strikingly, inhibiting Nodal signaling completely blocked re-aggregation.

**Figure 4.**
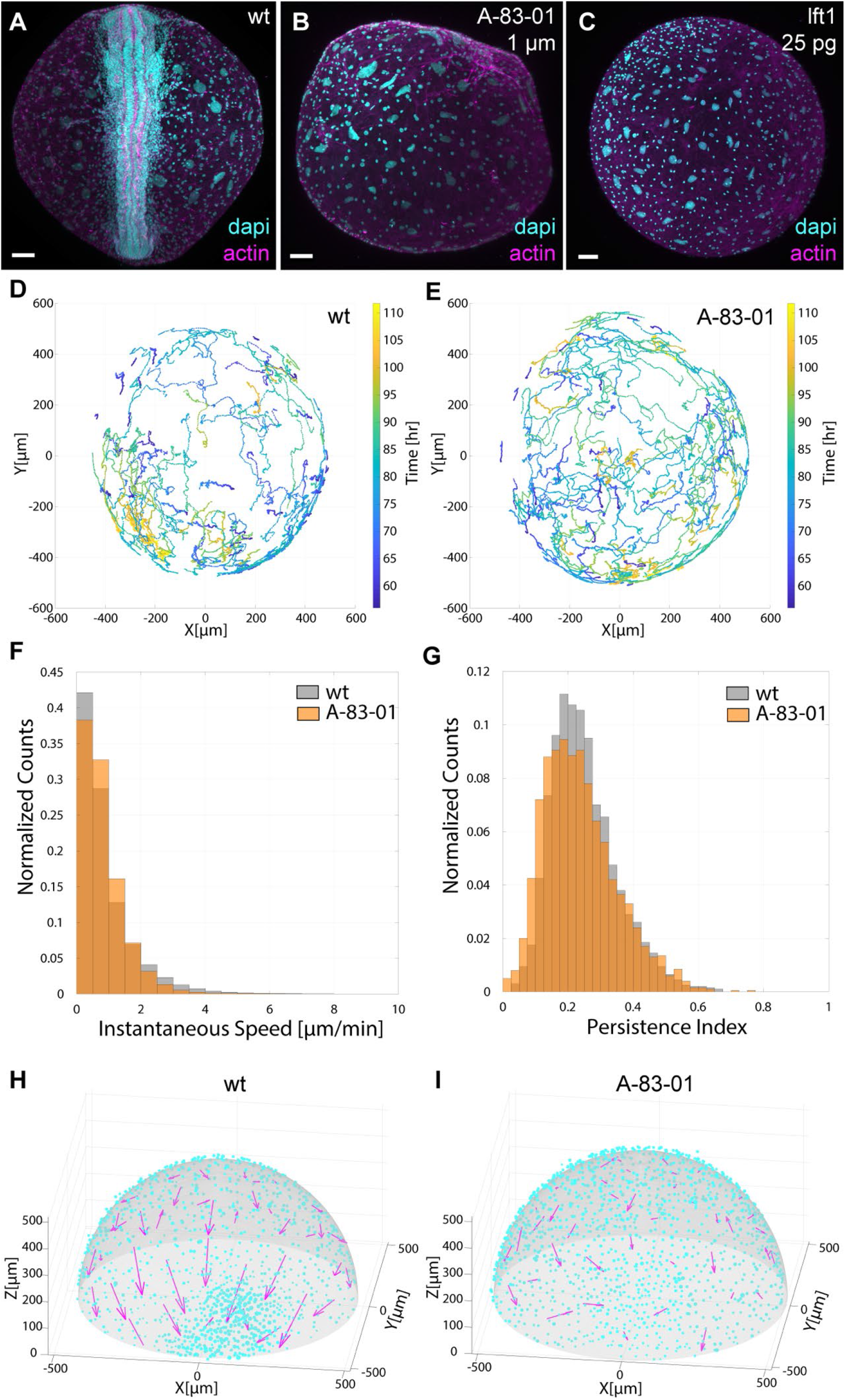
Nodal is required for coordinated morphogenesis. (**A**) Wild-type 10-somite stage embryo at day 5 (96.7% developed axes, n = 30). (**B**) Nodal inhibited embryo incubated with 1 μM A-83-01 from the dispersed phase to the incipient aggregate stage, then assayed at day 5 (0% developed axes, n = 43). (**C**) Embryo injected with 25 pg of *lefty1* mRNA at the 1-cell stage, then assayed at day 5 (5.9% developed axes, n = 17). (**D**) The top 200 cell trajectories with the most super-linear mean-squared displacement (MSD) curves from a time-lapse of wild-type development between 50-110 hpf. (**E)** The top 200 cell trajectories with the most super-linear MSD curves from a time-lapse of 2 μM A-83-01 inhibited development between 50-110 hpf. (**F**) Distributions of instantaneous speeds in wild-type and A-83-01 inhibited embryos. (**G**) Distributions of persistence indices in wild-type and A-83-01 inhibited embryos. (**H**) Comparison of local cell-cell alignment in wild-type and (**I**) A-83-01 inhibited embryos from 72-108 hpf. Magenta arrows represent the direction and magnitude of the local alignment in each area bin. Cyan dots indicate the position of each cell at 108 hrs. The aggregate is positioned at zero on the x-axis. Scale bars, 80 μm in (A) to (C).

To perform detailed phenotypic analysis of the aggregation defect caused by Nodal inhibition, we used light sheet microscopy from the dispersed phase until axis formation in transgenic *N. furzeri* expressing a nuclear fluorescent marker (movie S3 and S4). Using an automated 3D cell-tracking pipeline (*18*), we generated cell trajectories of individual blastomeres for wild-type and Nodal-inhibited embryos (Fig. 4, D and E, and fig. S5). By comparing cell behaviors (e.g. distributions of instantaneous speeds, persistence indices, and velocity autocorrelation decay times), we found that general motility was not affected by Nodal inhibition (Fig. 4, F and G, and fig. S5f). Next, we asked whether the local alignment and directionality of blastomeres was dependent on Nodal signaling. In the wild-type embryo, cell trajectories were on average more locally aligned, and mean cell directionalities pointed towards the aggregate (Fig. 4H). In contrast, cell trajectories were less aligned in the Nodal-inhibited embryo, and directionalities were random (Fig. 4I). These results reveal that Nodal signaling in *N. furzeri* does not affect the motility of individual deep cells but is important for coordinated migration and re-aggregation.

In summary, our results reveal three surprising divergences of early axis formation mechanisms in the annual killifish *N. furzeri* as compared to previously studied anamniotes. First, *huluwa* has devolved into a pseudogene and is inactive as a dorsalizing determinant. Second, embryonic axis formation only emerges when β-catenin stabilizes during re-aggregation. Third, Nodal signaling is required for coordinating cell migration during re-aggregation Thus, in the absence of Huluwa-dependent prepatterning, Nodal-coordinated migration allows re-aggregation and subsequent axis formation.

What are the conserved or divergent aspects of *N. furzeri* embryogenesis? Some of the later features of pattern formation appear conserved between *N. furzeri*, zebrafish and other fishes (*6, 19*). For example, *N. furzeri* β-catenin and *chordin* mark the dorsal organizer, and Nodal signaling and *tbx1* mark the presumptive mesoderm at the incipient aggregate stage. In contrast, at the blastula stage, *N. furzeri huluwa* is inactive as a dorsal determinant, and β-catenin does not accumulate asymmetrically. *N. furzeri* Nodal signaling has maintained its conserved role in the activation of organizer and mesoderm genes, but strikingly, it is also required for the directional migration and re-aggregation of the dispersed blastomeres. This early role of Nodal signaling differs from its established later roles in morphogenesis during gastrulation (*20–25*) and raises the question how Nodal signaling regulates individual cell migration.

Why does the annual killifish *N. furzeri* use such a divergent strategy of axis formation? In the wild, the dispersed phase is often prolonged by the entry into diapause I, a suspended state that allows survival during the dry season. There are three possibilities why this dispersed state might be incompatible with prepatterning. First, it is conceivable that during diapause, cellular damage could accumulate and affect a prepatterned dorsal axis. Dispersion might serve as a buffer from this type of damage, allowing cells to re-aggregate using the remaining uncommitted cells (*4, 26, 27*). Second, the cellular rearrangements that occur during dispersion might erase maternal prepatterns due to the loss of prior spatial information. Indeed, dissociation and aggregation of zebrafish blastomeres can impede axis formation (*28*). Third, prolonged developmental arrest could flatten previously formed morphogen gradients (*29*). Thus, dispersion might facilitate survival during diapause but may preclude maternal prepatterning as a viable mechanism for axis formation.

What is the relationship between the apparent self-organization of the *N. furzeri* aggregate and the embryonic patterning of other metazoans? Remarkably, human and mouse embryonic stem cells can be induced to self-organize, express germ layer markers and resemble gastrula stage embryos (*30*). Moreover, even species that normally prepattern their axes (e.g. sponges, anemone, and amphibians) have been shown to have the capacity to self-organize after experimental dissociation (*31–33*). These observations suggest that when challenged, some animals that normally prepattern their embryos may have the ability to pattern their axes through an alternative, self-organizing route. We speculate that in annual killifish, this self-organized route of development may have become fixed in order to survive the selective pressures imposed by their extreme environment.

## Acknowledgments

We thank the following individuals: Nathan Lord, Adam Carte, Annika Nichols and Yinan Wan for helpful discussions and critical reading of the manuscript; Kate McDole for providing advice about automated cell tracking; Chi-Kuo Hu and Itamar Harel for training and providing embryos; Brittany Hughes, Jessica Miller, Karen Hurley, Kathryn Berg, and Tomoki Kimura for animal husbandry; Douglas Richardson of the Harvard Center for Biological Imaging for microscopy support; and Joel Sohn for helpful discussions.

## Author contributions

P.B.A, D.C.A, and A.F.S conceived the study and wrote the paper. P.B.A. performed all experiments. D.C.A. analyzed the light sheet data.

## Supplementary Materials

Materials and Methods

Figures S1-S5

Movies S1-S4

References (34-41)

