## Supplementary material for "Axis formation in annual killifish: Nodal coordinates morphogenesis in absence of Huluwa prepatterning": Methods and supplementary materials

### Materials and Methods:

#### Animal husbandry and stable transgenesis

An inbred strain of *Nothobranchius furzeri*, originating from Gonarezhou National Park, Zimbabwe was used in each experiment. Animal husbandry was performed as previously described (*8*). The protocol was modified to be peat-free, using 0.5 mm glass disruptor beads (USA Scientific, 7400-2405) as egg-laying material and damp, 90 mm filter paper (Whatman, 1001-090) for egg incubation.

For transgenesis two stable lines were generated using the Tol2 system as previously described (*34*). The β-catenin nanobody (BC1:egfp) was subcloned from a previous study (*12*) into the Tol2 vector and driven by the 1316 bp *β-actin* promoter. The nuclear fluorescent marker used in the light sheet experiments is a human H2B Dendra2 fusion driven by the 1316 bp *β-actin* promoter. Both transgenic lines are available upon request.

#### Embryo preparation and imaging

*N. furzeri* embryos were collected from breeding adults and injected at the 1-cell stage with approximately 25 pg of mRNA. Embryos were incubated in Activin A Receptor Type 1B and Activin A Receptor Type 1C inhibitor A-83-01 (Sigma, SML0788) at a concentration of 1-2 μM for the durations indicated in the figure legends. Embryos were raised at 28°C and fixed at the appropriate stage overnight in 4% paraformaldehyde (EMS, 15714-S) at 4°C. The next day the tissue was rinsed in 0.05% Tween-20 in PBS and the chorions were manually removed using fine forceps. The tissue was then cleared in a series of four 20-minute washes of 0.05% Tween-20 in PBS. Actin and DNA were stained overnight with Alexa-647 conjugated phalloidin (Fisher, A22287) at a dilution of 1:100 and DAPI (Fisher, D1306) as recommended by the manufacture, respectively.

For antibody staining, tissues were fixed overnight in 4% paraformaldehyde (EMS, 15714-S) at 4°C and quickly washed in 0.05% Tween-20 in PBS. Chorions were manually removed and washed four times in 0.05% Tween-20 in PBS for 20 minutes. Embryos were blocked using 1% normal goat serum and 1% DMSO for one hour. Afterward samples were incubated with the mouse anti-E-Cadherin (BD, 610181), rabbit anti-β-catenin (abcam, 209860), or rabbit anti-Smad2/3 (Cell Signaling, D7G7) antibody at a dilution of 1:500 overnight at 4°C. The following day the tissue was washed four times in 0.05% Tween-20 in PBS for 20 minutes. Then the samples were incubated with the appropriate Alexa Fluor secondary antibody at a dilution of 1:500 in 1% DMSO for two hours at room temperature, and finally washed in 0.05% Tween-20 in PBS, four times for 20 minutes each.

For confocal imaging samples were mounted in 1.2% low melt agarose in PBS in plastic 60 x 15 mm petri dishes. Confocal images were acquired on an LSM upright 880 Zeiss microscope using a W plan-apochromat 10x or 20x objective. Confocal stacks were typically 150-300 μm thick. Images were rendered in 3D using Bitplane Imaris software. Time-lapse microscopy experiments were acquired on a Zeiss light sheet Z.1 at intervals of 150 seconds from a single view. Embryos were mounted in capillaries containing 1.2% low melt agarose in filtered aquarium water. The light sheet chamber was filled with filtered aquarium water containing 100 units/mL of penicillin-streptomycin (Gibco, 15140122) to prevent bacterial growth. Brightfield images were taken with an LG G6 camera phone with a Zeiss Stemi 2000.

Experimental perturbations were performed in triplicate and at least two biological replicates were quantified. The scored replicates were totaled and are shown in each corresponding figure legend. Unfertilized embryos were excluded from the analysis. No randomization or blinding was performed on samples.

**Molecular biology for misexpression studies**

*N. furzeri* genomes from the Fritz Lipmann Institute and Stanford were used to identify coding transcripts (*9*, *35*). Coding sequences were PCR amplified and cloned into a pCS2 vector using *Xho1/Xba1* restriction site for *lefty1* (5'-ATGGATGTCCTTGGCGCTTGCCTC and 3'-TCATACCAGGGACATGTCCATC), and *EcoR1/Xho1* for *N. furzeri huluwa* (5'-ATGGAAGCAGAACACATGAGTC and 3'-TTAGACCCAGTACTGCTTTGTG) from cDNA made from cleavage stage *N. furzeri* embryos. *D. rerio huluwa* (XM_698288.8), and *K. marmoratus huluwa* (XM_025004129.1) were synthesized by Twist Biosciences and cloned in the pCS2 expression vector.

***In situ* hybridization**

The fluorescent *in situ* hybridizations were performed as described using a method for *D. rerio* (*36*). *N furzeri.* specific mRNA probes were synthesized using linearized cDNA templates for *ndr2* (5'-ATGCGCTCTTTTGGAGCTCCGGGAG and 3'-TCATTGGCAGCCACACTCTTCCACG), *lft1* (5'-ATGGATGTCCTTGGCGCTTGCCTC and 3'-TCATACCAGGGACATGTCCATC), *chrd* (5'-GCGTTTTAAAGGACCTGAGTGTTG and 3'-CAGGACAGCAGGAATTCTCCGTACG), *tbx1* (5'-GACATTAAACTGACCCTGGAGGA and 3'-GAAGAAGACCCTGAAGGTGTGAT), *sox2* (5'-ATGTATAACATGATGGAGACTG and 3'-TCACATGTGTGTTAACGGCAGCG), *otx2* (5'-GGAAACAACGACGAGAAAGGAC and 3'-CCAAGCAATCGGCATTGAAGTTC), and *gbx1* (5'-ATGCAACGACCAGGCGGGCAG and 3'-TCATGGTCTTGTCCCCTGCTCTAT), which were made using the StrataClone PCR Cloning Kit (Agilent Technologies, #240205).

**Phylogenetic analysis**

Orthologous Huluwa coding sequences were downloaded from NCBI and aligned using the MUSCLE algorithm (*37*) (*Python bivittatus*, XP_025031808, *Xenopus laevis* XP_018122910, *Danio rerio* XP_703380, *Astyanax mexicanus* XP_007256672, *Salmo salar* XP_013983523, *Oryzias latipes* XP_011479863, *Haplochromis burtoni* XP_005917435, *Larimichthys crocea* XP_019124609, *Takifugu rubripes* XP_011602919, *Poecilia mexicana* XP_014834893, *Xiphophorus couchianus* XP_027882250, *Kryptolebias marmoratus* XP_024859897, and *Nothobranchius furzeri* GAIB01045173). Phylogenetic analysis was performed using PhyML 3.0 (*38*) using Geneious Prime. The transmembrane domains were predicted using Transmembrane Hidden Markov models.

### Automated cell tracking

Background subtraction was performed on all acquired raw images using custom code written in MATLAB. Briefly, a kernel fit was performed on the histogram of all pixels for a given 3D stack for each time point, and the peak was set as the background, with intensities below the peak being set to 0. Background-subtracted images were converted into the KLB format, and the TGMM pipeline was used to segment and track cells (*18*), using a background thresholds of 150 and 100, and tau level of 12 for the wild-type development and Nodal inhibition time-lapses, respectively. Tracks were imported into ImageJ via the MaMuT plugin (*39*) for manual validation, and validated tracks were imported into MATLAB using custom code.

### Rotational drift correction

To correct for rotational drift of the embryo inside the chorion, we wrote custom MATLAB code. Briefly, imported tracks were centered at the origin via a sphere fit to all cell positions, with a maximum distance constraint of 5 pixels. The displacement of each cell between two consecutive time points was then described via Euler angles around the origin. All Euler angles for a given time point were averaged to estimate the mean rotation of the sample at that time point; we assume that averaging eliminates any variation due to true cell movement, leaving behind coordinated rotation of all cells due to drift between consecutive time points.

The mean Euler angles were converted to rotation matrices, and the transpose of each matrix was stored as a Rotational Drift Correction Matrix (*RDCM*) for each time point. We decided to choose the first time point of each time-lapse as the reference orientation, thus defining:

$$RDCM(t = 0) = I$$

for each cell *i* at time point *t*, the 3D position vector $x_i^t$ was corrected as:

$$x_{i,corr}(t) = \prod_{\tau=0}^{t} RDCM(\tau) * x_i(t)$$

### Track filtering

Automated tracking via TGMM is error prone, often leading to very short tracks, or erroneous tracks with large spatial leaps between consecutive steps (often due to incorrectly linked cells). To filter for high quality tracks, we removed any tracks generated by TGMM that (a) tracked objects for less than 100 frames, and (b) had any leaps between consecutive time points 7x larger than the nucleus of the object. These filters were set heuristically. This reduced the total number of tracks to 9,689 for the wild-type development time-lapse, and 6,006 for the Nodal inhibition time-lapse.

### Mean-squared displacement (MSD) analysis and track properties

MSD curves for all filtered tracks were calculated using msdanalyzer (*40*). A linear fit to the first 25% of the MSD curve of each tracks in log-log space was then performed. The slope of this fit corresponds to the exponent of time in the time-MSD curve, noted by α. This exponent is indicative of the type of motion the cell follows: for a random walking cell, the MSD was expected to be linear with time, leading to $\alpha = 1$. For a cell moving super-diffusively, the MSD grows faster than linear, leading to $\alpha > 1$, and for a confined cell, the MSD grows slower than linear, leading to $\alpha < 1$. Trajectories with varying α values, as well as their corresponding MSD curves are shown in fig. S5, A and B.

Tracking resulted in trajectories of both EVL cells and deep blastomeres. However, the aggregation behavior is driven solely by the blastomeres. Since EVL nuclei are confined in an epithelium, these tracks were assumed to have lower α values. Therefore, to study isolated blastomeres, the top 2,000 tracks with the largest α values were chosen, and these tracks from both time-lapses were used for further analysis. Persistence indices of tracks were calculated by taking the ratio of the net displacement of trajectories by the integrated displacement across all time points. Velocity autocorrelation was calculated using msdanalyzer (*40*).

**Local cell-cell alignment analysis**

To quantify the coordination of cell trajectories over the time-lapses, a vector version of the scalar order parameter introduced by Vicsek *et al.* (*41*) was used. This parameter describes the alignment of unit velocity vectors in a dataset, and is given by the equation:

$$\vec{\Phi}(t) = \frac{1}{N}\sum_{i=1}^{N}\frac{v_i(t)}{|v_i(t)|}$$

Where $v_i$ is the velocity vector of cell *i* at a given time point *t*, and *N* is the number of cells. The magnitude of $\vec{\Phi}$ is a measure of the alignment of cell directions, while the direction of $\vec{\Phi}$ indicates the mean directionality of cells considered in the average.

There are two issues with the use of a global alignment parameter for cell trajectories given the data: First, cell trajectories are confined on a sphere-like 2D manifold set by the yolk, which restricts the observable directions of cells depending on their position on the embryo. Second, since cells re-aggregate towards a single point, the directionalities of cells approaching from opposing sides would cancel, resulting in a seemingly reduced global alignment parameter, despite potential high local alignment of trajectories.

To address these issues, this parameter was evaluated locally along the surface of the sphere-like embryo. First, to generate approximately equal-area bins along the embryo surface, a triangular icosphere mesh with the mean radius of the embryo was used. To determine the appropriate length scale for the mesh, the magnitude of the mean alignment of cells was calculated as a function of distance from a given cell and averaged this across all cells and time points. Then, an exponential decay was fit to the resulting graph to determine a mean decay length scale for cell-cell alignment. For the wild-type development time-lapse, this was 197.8 μm, and for the Nodal inhibition time-lapse, it was 176.7 μm. These length scales were matched approximately to the mesh dimension size for each embryo, using the following Matlab suite: https://www.github.com/AntonSemechko/S2-Sampling-Toolbox.

To calculate a local alignment parameter, cells were binned by assigning them to the closest triangle on the mesh. The alignment parameter $\vec{\Phi}$ was calculated for each bin at every time point. For Figure 4g, i $\vec{\Phi}$ values were averaged for each bin over the 72-108 hr time window to get mean local alignment values during the aggregation stage.

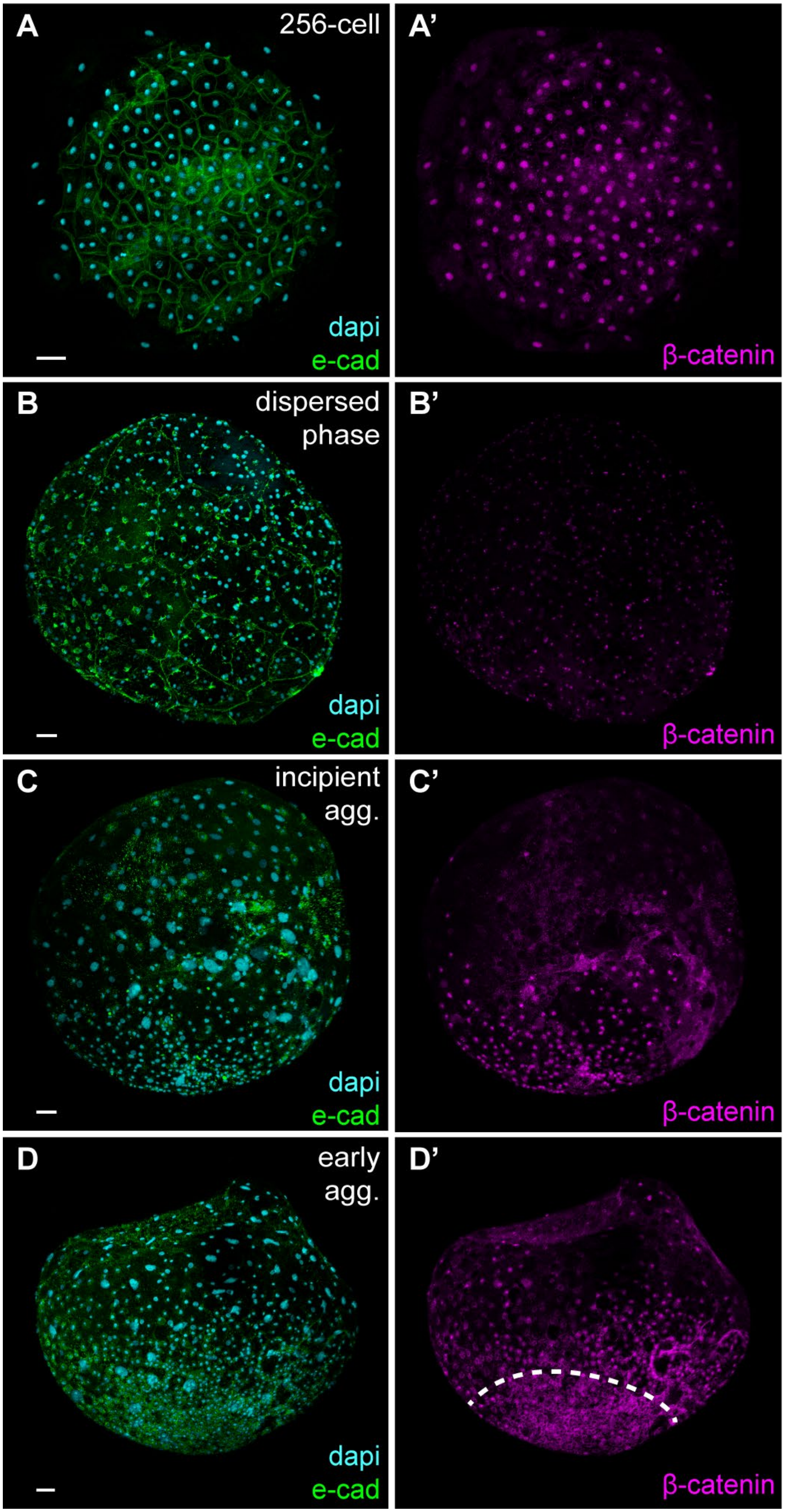
A
256-cell
dapi
e-cad
A'
β-catenin
B
dispersed
phase
dapi
e-cad
B'
β-catenin
C
incipient
agg.
dapi
e-cad
C'
β-catenin
D
early
agg.
dapi
e-cad
D'
β-catenin

**Fig. S1. Anti-β-catenin staining during early *N. furzeri* development.** (**A**) Animal view of 256-cell stage embryo prior to dispersion. (**A'**) Nuclear β-catenin is seen in all cells. (**B**) Lateral view of dispersed phase embryo at the end of epiboly. (**B'**) β-catenin signal is diminished compared to pre-dispersion stage embryos. (**C**) Lateral view of an incipient aggregate stage embryo. (**C'**) Nuclear β-catenin is seen in the bottom hemisphere of the embryo in re-aggregating cells. (**D**) Lateral view of early aggregate stage embryo. (**D'**) Nuclear β-catenin is enriched in cells at the periphery of the aggregate, above the dotted line. Scale bars, 50 μm.

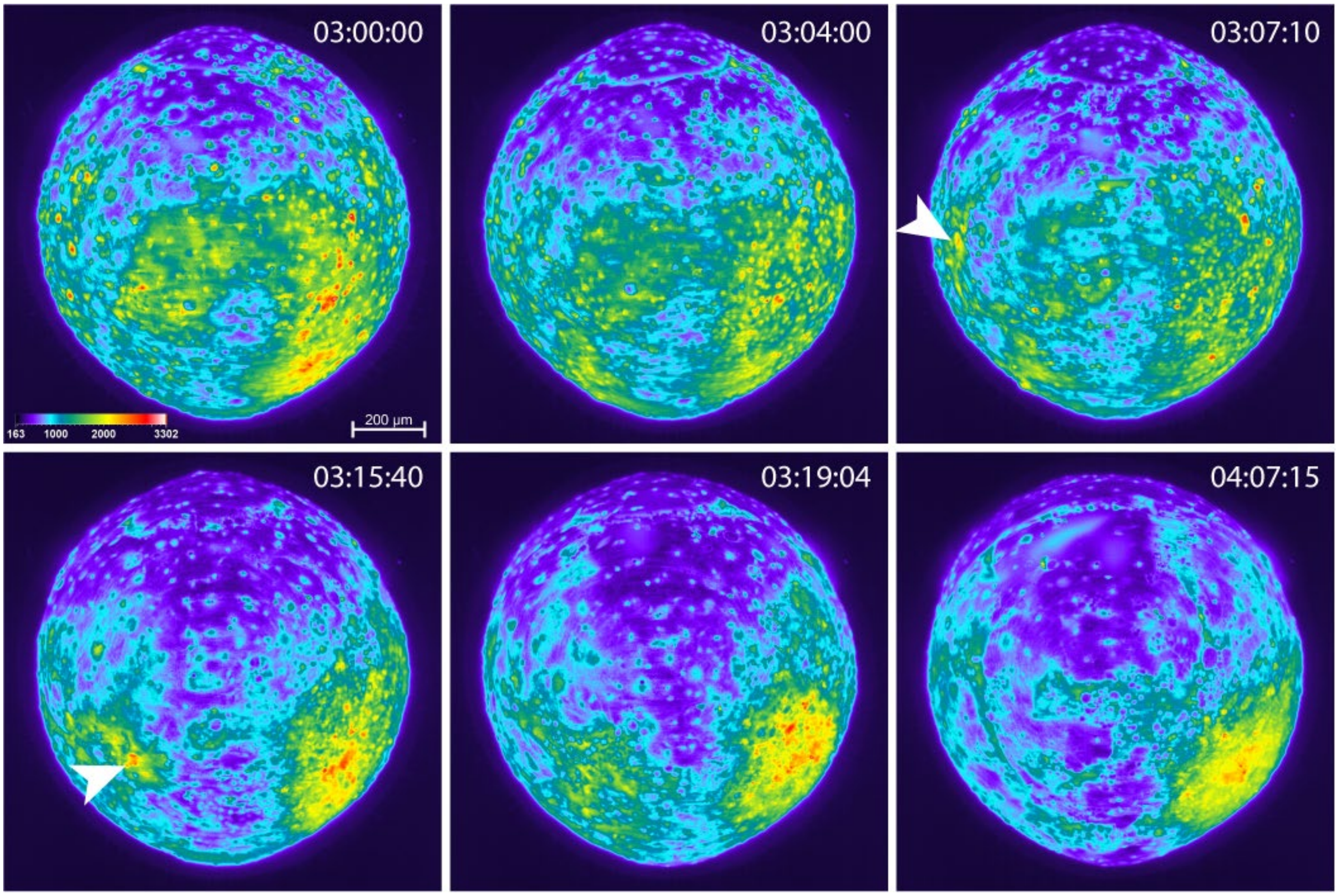


**Fig. S2. Nanobody reveals β-catenin dynamics during re-aggregation.** Snapshots from a time-lapse of transgenic *N. furzeri* embryo that expresses the nanobody BC1:egfp, starting from incipient aggregation stage (72 hpf). Timestamp indicates days: hours: minutes. The signal intensity is color coded (see intensity scale). The aggregate forms on the right lower quadrant (yellow, red signal). Local transient increases in nuclear β-catenin occurred near the aggregate (white arrowheads). The full dynamics can be seen in movie S2.

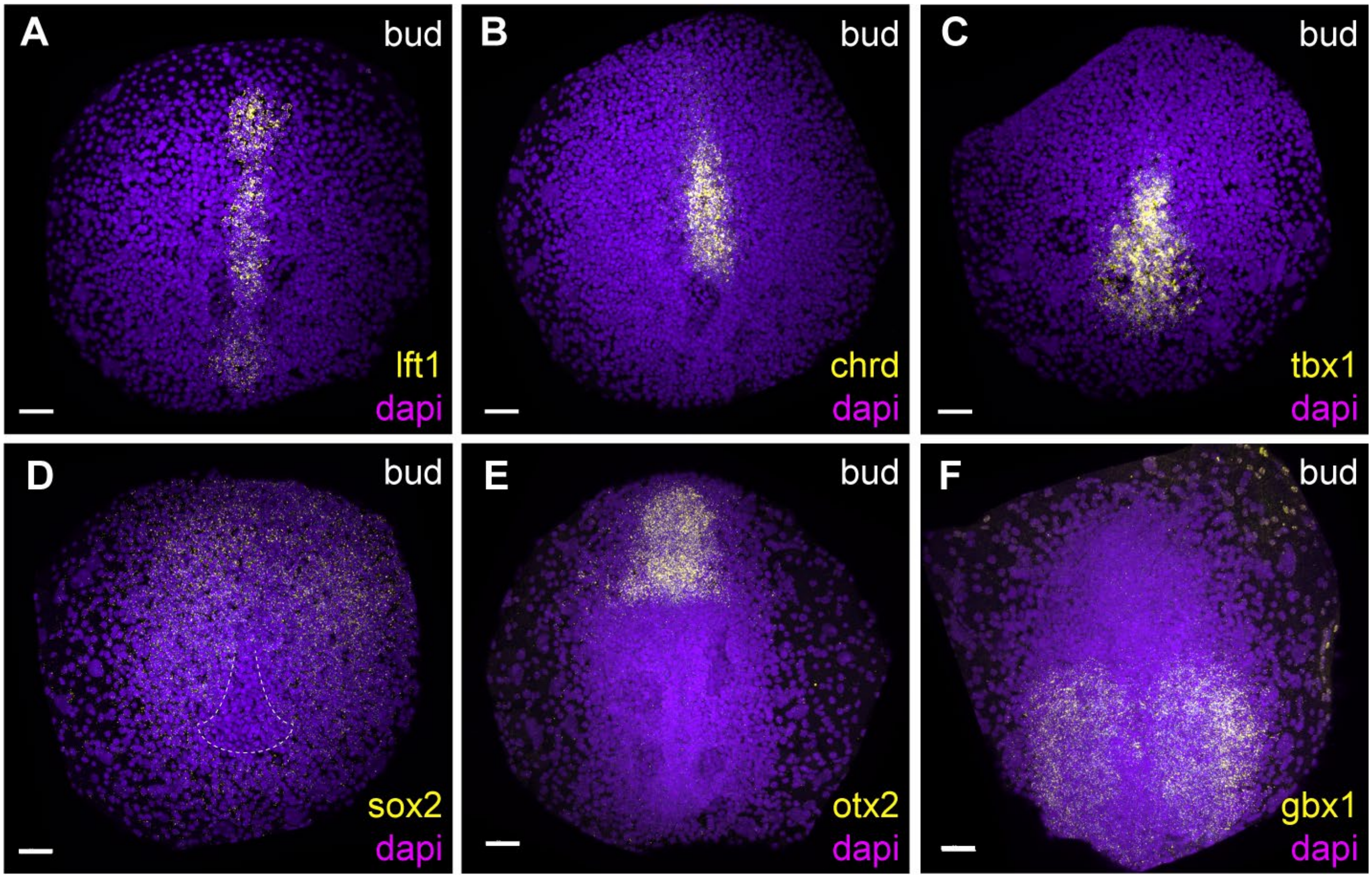


**Fig. S3. Expression of Nodal targets and other key marker genes during the bud stage.** (**A**) *Lefty1* mRNA expression. (**B**) *Chordin* mRNA expression. (**C**) *Tbx1* mRNA expression. (**D**) *Sox2* mRNA expression. Dotted line triangle demarcates the involuting mesoderm; see *tbx1* mRNA expression in (C), where there is no expression of *sox2*. (**E**) *Otx2* mRNA expression. (**F**) *Gbx1* mRNA expression. In all images anterior is up. Scale bars, 50 µm.

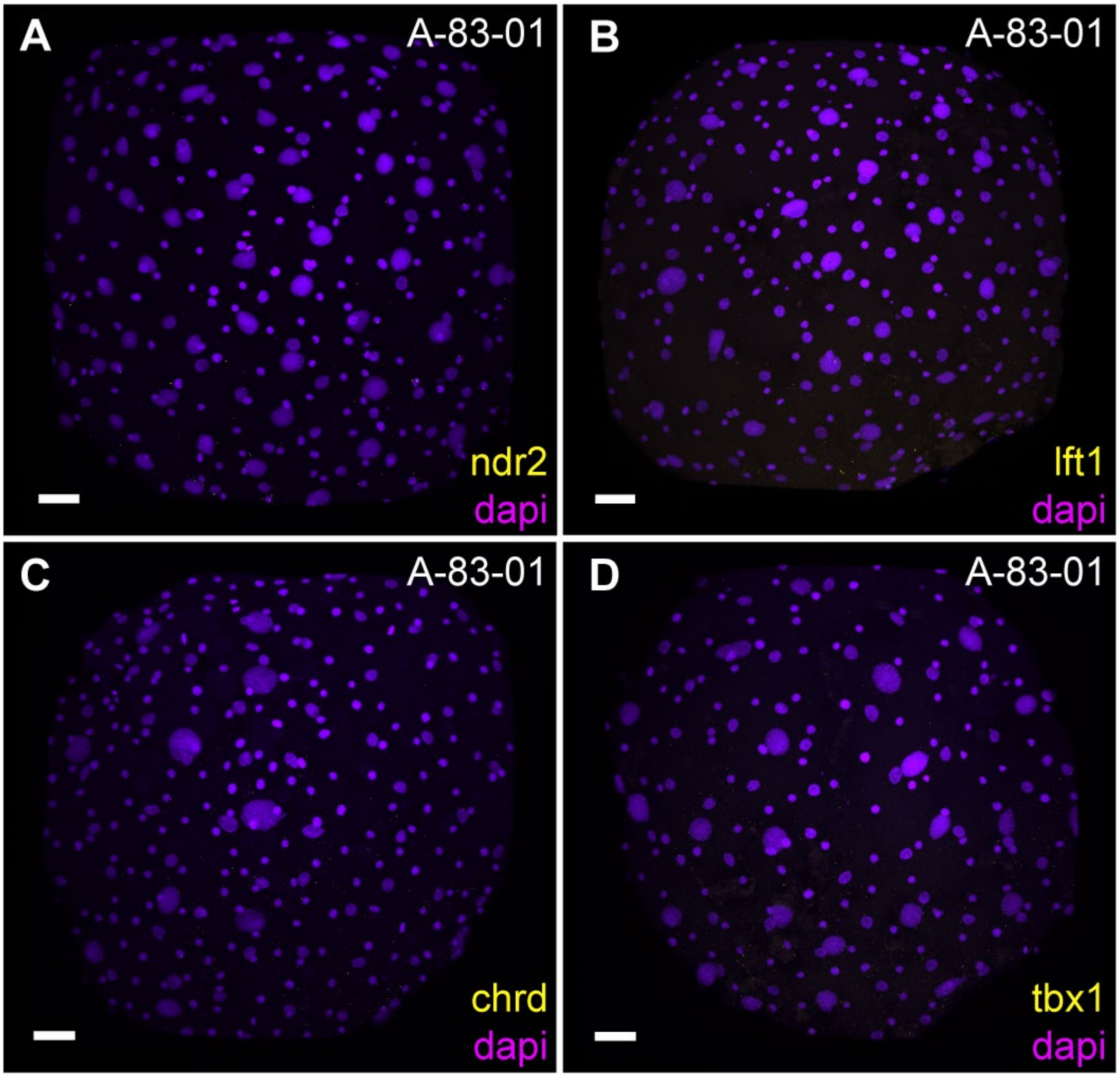


**Fig. S4. Nodal inhibitor blocks target gene expression.** (**A**) *Ndr2* mRNA expression assayed at the incipient aggregate stage after incubation with 1 μM A-83-01, starting from the dispersed phase. (**B**) *Lefty1* mRNA expression assayed at the incipient aggregate stage after incubation with 1 μM A-83-01, starting from the dispersed phase. (**C**) *Chordin* mRNA expression assayed at the incipient aggregate stage after incubation with 1 μM A-83-01, starting from the dispersed phase. (**D**) *Tbx1* mRNA expression assayed at the incipient aggregate stage after incubation with 1 μM A-83-01, starting from the dispersed phase. Scale bars, 50 μm.

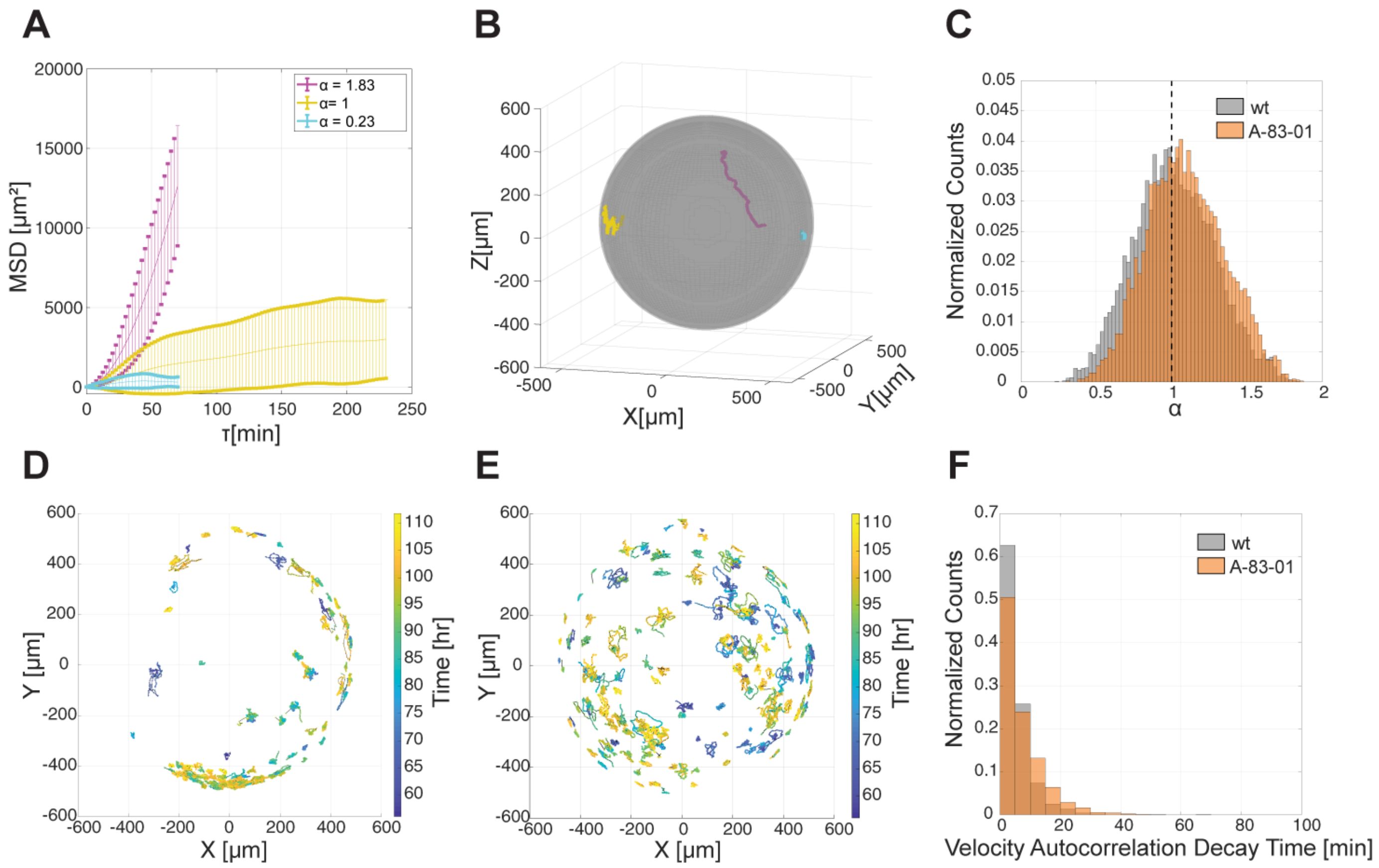


**Fig. S5. Track sorting and characterization**. (**A**) Examples of mean-squared displacement (MSD) curves calculated for tracks from the wild-type development dataset. τ indicates time lag, and α indicates the exponent of τ in an exponential fit to each curve. Error bars indicate standard deviation. (**B**) Trajectories from (A) plotted onto a sphere with the mean radius of the wild-type developing embryo. (**C**) Normalized distributions of MSD fit exponents, or α values, for the wild-type development and Nodal inhibition time-lapses. Dotted line marks α = 1. (**D**) 200 tracks with the lowest α values from wild-type development time-lapse and (**E**) Nodal inhibition time-lapse plotted in the X-Y plane. (**F**) Normalized distributions of the velocity autocorrelation decay time for tracks in the wild-type development and Nodal inhibition time-lapses.

**Movie S1**
This movie shows the lateral view of a transgenic *N. furzeri* embryo expressing BC1:egf during dispersion. The time lapse is approximately 28 hours and spans from the beginning of the dispersed phase to the end of epiboly (QuickTime; 22.3 MB).

**Movie S2**
This movie shows the lateral view of a transgenic *N. furzeri* embryo expressing BC1:egf during aggregation. The time lapse is approximately 26 hours and spans from the incipient aggregate stage to the early aggregate stage (QuickTime; 20.9 MB).

**Movie S3**
This movie shows the lateral view of a wild-type transgenic *N. furzeri* embryo expressing H2B:Dendra2. The time lapse is approximately 60 hours and spans from the end of epiboly to the bud stage. The embryo aggregates near the bottom of the egg (MPEG; 23.5 MB).

**Movie S4**
This movie shows the lateral view of a Nodal-inhibited transgenic *N. furzeri* embryo expressing H2B:Dendra2. The embryo was incubated with 2 μM A-83-01. The time lapse is approximately 60 hours and begins at the end of epiboly (MPEG; 23.6 MB).
